# Invasive slipper limpets *Crepidula fornicata* act like a sink, rather than source, of *Vibrio* spp.

**DOI:** 10.1101/2021.12.16.472931

**Authors:** Emma A. Quinn, Sophie H. Malkin, Jessica E. Thomas, Ryan Poole, Charlotte E. Davies, Andrew F. Rowley, Christopher J. Coates

## Abstract

A large knowledge gap exists regarding the disease profile and pathologic condition of the invasive, non-native, slipper limpet *Crepidula fornicata*. To help address this, we performed a yearlong health survey across two sites in South Wales UK – subtidal Swansea Bay and intertidal Milford Haven. In total, 1,800 limpets were screened systematically for haemolymph bacterial burdens using both general and vibrio-selective growth media (TSA +2% NaCl and TCBS, respectively), haemolymph (blood) inspection using microscopy, a PCR-based assay targeting *Vibrio* spp., and multi-tissue histology. Over 99% of haemolymph samples contained cultivable bacterial colony forming units, and 83% of limpets tested positive for the presence of vibrios *via* PCR (confirmed *via* Sanger sequencing). Vibrio presence did not vary greatly across sites, yet a strong temporal (seasonal) effect was observed – significantly higher bacterial loads during the summer. Binomial logistic regression models revealed larger (older) limpets were more likely to harbour vibrios, and the growth of bacteria on TCBS was a key predictor for PCR-based vibrio detection. Histological assessment of >340 animals revealed little evidence of inflammation, sepsis, or immune reactivity despite the gross bacterial numbers. We contend that slipper limpets are not susceptible to bacteriosis at either site surveyed, or do not to harbour vibrios known to be pathogenic to humans. The lack of susceptibility to local pathogenic bacteria may explain, in part, the invasion success of *C. fornicata* across this region.

## Introduction

Molluscs form important economic and ecological components of marine ecosystems (FAO, 2017; Sowa et al., 2019). They aid in determining benthic community structure, nutrient cycling, act as food sources for higher trophic levels, stabilize shorelines, and maintain water quality alongside their exploitation for commercial aquaculture and wild-harvest industries (Goss-Custard et al., 2004; Jansen et al., 2012; Kellogg et al., 2014; Tomatsuri and Kon, 2017; Walles et al., 2015). As many aquatic molluscs filter-feed, this can lead to the accumulation of diverse bacterial consortia, notably *Vibrio* spp., with some hosts acting as key reservoirs/sources of pathogenic ecotypes. The Atlantic slipper limpet *Crepidula fornicata* is one such filter feeder, and represents a prolific invasive, non-native species (INNS) introduced to Europe in the 1870s with shipments of American oyster (*Crassostrea virginica*; Blanchard, 1997). For over a century, *C. fornicata* has occupied European coastlines reaching >4700 limpets m^-2^ *in extremis* (Marennes-Oléron, France; de Montaudouin and Sauriau 1999). The first report of *C. fornicata* in Welsh shores came from Milford Haven, Pembrokeshire in the 1950s (Cole and Baird, 1953), and over the following decade established itself throughout the waterway (e.g., Hazelbeach; Crothers, 1966). These limpets have gained notoriety as an “oyster pest” due to their ability to modulate benthic community composition, sedimentation and pseudofaeces accumulation, food and trophic competition with native and Pacific oysters (*Ostrea edulis* and *Crassostrea gigas*), queen scallops (*Aequipecten opercularis*) and blue mussels (*Mytilus edulis*; Thieltges et al., 2003 and 2006; Thieltges, 2005; Kriefall et al. 2018; Richard et al., 2006; Preston et al., 2020). There are, however, conflicting reports on the nature o*f C. fornicata’s* adverse influence. De Montaudouin et al. (1999) demonstrated in laboratory trials that *C. gigas* growth, macrozoobenthic density and diversity was not negatively impacted by the presence of *C. fornicata*. Using natural history records, Hayer et al. (2019) reconstructed the changes in distribution and diversity over the last 200 years in the North Sea – concluding that the decline of *O. edulis* was near completion before the introduction of *C. fornicata*. Conversely, Le Pape et al. (2004) showed that *C. fornicata* presence can reduce the density of young-of-the-year sole (*Solea solea*) in flatfish nurseries by limiting their ability to bury into the sediment, and Thieltges (2005) concluded that *C. fornicata* is an important mortality factor for *M. edulis in situ*.

There is much interest in *C. fornicata* as an ecosystem engineer and a driver of ecological phase shift outside its native range, yet there is a paucity of knowledge with respect to its pathogen/parasite profile, immunobiology and epizootiology (Quinn et al., 2020). Alternatively, this species may be resistant to many common disease-causing agents. Such data are key to considering the risk of disease transfer to other co-located animals like commercially sensitive oysters (*C. gigas*) and mussels *(M. edulis*). There is some evidence that *C. fornicata* can act as a sink of trematode parasites that use mytilids as intermediate hosts, but the limpets themselves did not appear susceptible (Pechenik et al., 2001; Thieltges et al., 2006). Despite an exhaustive literature search, we found one published study with respect to a systematic disease survey of *C. fornicata*. Le Cam and Viard (2011) monitored the temporal prevalence of a boring sponge, *Cliona celata*, in a French population of *C. fornicata* over three years (2004-2007). They examined a total of 12,049 individuals and found that on average 43% were infested by the sponge (skewed toward females) although this infestation appeared to have limited effects on the host. With respect to bacterial presence, Choquet (2004) isolated a strain of *Vibrio tapetis* – the cause of brown ring disease in clams – from *C. fornicata*.

Members of the bacterial genus *Vibrio* blight marine invertebrates – highly virulent strains are associated with mass die-off events of commercial bivalves (e.g., *Vibrio aestuarianus* in Pacific oysters; Labreuche et al., 2006; Lupo et al., 2019) and crustaceans (e.g., *Vibrio parahaemolyticus* causes acute hepatopancreatic necrosis disease in penaeid shrimp; Kumar et al., 2020). Vibrios are Gram-negative, curved rods of the class *gamma-Proteobacteria* (Williams et al., 2010), which comprise over 100 species grouped into 14 clades (Romalde et al., 2014). Vibrios are almost ubiquitous in aquatic environments, found naturally in marine, estuarine, and freshwater. They can survive in a diverse range of habitats, having the ability to colonize fish and other marine invertebrates, associate with plankton and algae, and having the capacity to form biofilms on biotic and abiotic surfaces (Reen et al., 2006). Many *Vibrio* species are harmless, yet some can cause disease in both humans and invertebrates (Froelich and Noble, 2016) with ∼12 *Vibrio* spp. that cause illness in humans – usually through the ingestion of contaminated food. The major agents of shellfish poisoning are *Vibrio parahaemolyticus, V. vulnificus*, and *V. cholerae*, leading to gastroenteritis, wound infections, and septicaemia. *V. parahaemolyticus* is responsible for the highest incidences of seafood-associated bacterial gastroenteritis in several countries, e.g., United States of America and Asia (Liu et al., 2004; Scallan et al., 2011).

Various diagnostic techniques can be used to detect vibrios. One standard method is through microbiological enrichment, namely thiosulfate-citrate-bile salts agar (TCBS), used routinely as the selective medium for vibrios. *Staphylococcus, Flavobacterium, Pseudoalteromonas, Aeromonas* and *Shewanella* isolates can also grow on TCBS, as such, the medium is not entirely restricted to vibrios. Vibrio strains that can utilise sucrose form yellow colonies on the medium, whereas the others are green (Thompson et al., 2004). Molecular methods such as PCR are also used to detect the presence of *Vibrio* (Vezzulli et al., 2012) – a combination of the two approaches is ideal.

Our aim was to perform the first large-scale temporal survey for vibrio-like bacteria in the alien invader, *C. fornicata*, across two affected locations in Wales (U.K.) – Swansea Bay (subtidal sampling within a native oyster restoration zone) and Milford Haven (intertidal sampling among an area of key fisheries/culture industries). To achieve this, we sampled 75 limpets per site per month for one year (January-December 2019) and used a multi-resource screen for determining vibrio presence. Liquid tissue (haemolymph) was studied for immune cell (haemocyte) numbers, plated on general (TSA +2% NaCl) and selective (TCBS) culture media for bacterial load quantification, and processed for DNA extraction and PCR-based diagnosis (universal vibrio primers; Vezzulli et al., 2012). Solid tissues – including the digestive gland and gills – were examined using H&E histology.

## Materials and Methods

### Sampling regime and site descriptions

Sampling efforts for both intertidal and subtidal collection of live adult *C. fornicata* (n = 1,800) took place at two sites in South Wales that are known to have well established populations (**Fig. 1a**). Swansea Bay (n = 900; 51.570345, -3.974591) was sampled subtidally *via* dredging, whereas limpets at Milford Haven (Hazelbeach; n = 900; 51.7042, -4.971295) were handpicked intertidally. Sample size calculations using an α-value of 0.05 and desired power >80% indicated that a minimum of 57 (1-sided test) up to 73 (2-sided test) limpets were required based on an *a priori* prediction of 15% (p1) to 35% (p2) prevalence of ‘diseased’ animals (in line with observations made by Le Cam and Viard (2011) when screening limpets for *C. celata*).

**Fig. 1.**
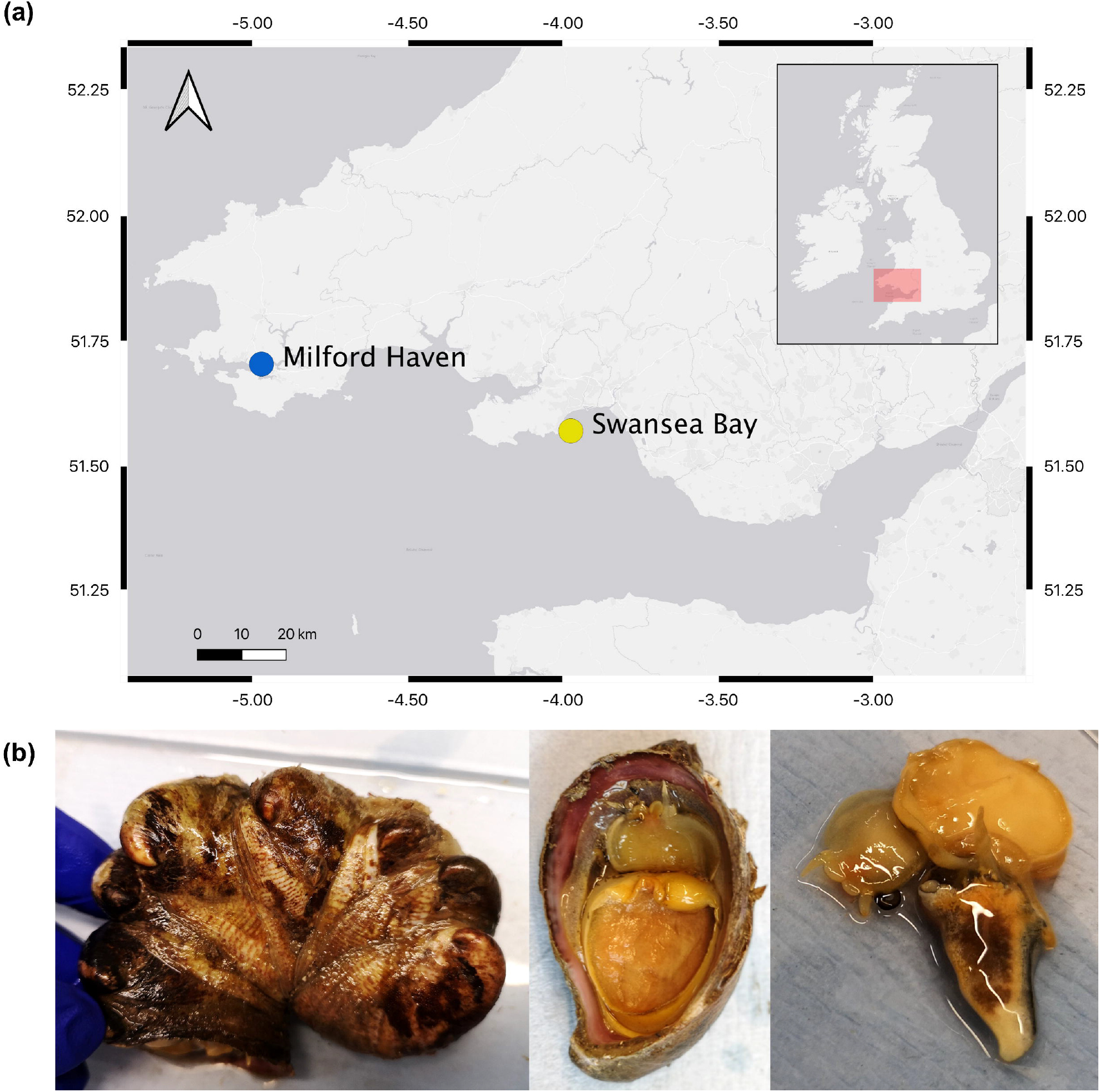
Collection and processing of slipper limpets (*Crepidula fornicata*). **(a)** Geographic locations of both sample sites in South Wales, UK. Maps were created in QGIS 3.10.3-A Coruña. Inset, geographic location of Swansea Bay within the United Kingdom (red box). **(b)** Each limpet is separated from the stack and a blunt ended probe is used to detach tissue mass/flesh from the aperture, while the haemolymph pools in the shell.

Swansea Bay is a shallow embayment with depth generally <20 m Ordnance Datum (OD), located on the northern coastline of the Bristol Channel (Pye and Blott, 2014). The bay stretches ∼12 km from Mumbles Head to Port Talbot. Swansea Bay’s tidal range is one of the largest in North-West Europe with a mean spring tide of 8.6 m and mean neap tide of 4.1 m (Collins et al., 1979; Shackley and Collins, 1984). Sediments are mostly fine and medium sand in the inner bay with an increasing proportion of mud close inshore to the west, due to protection from wave exposure by Mumbles Head, and shallower waters slowing tidal currents (Smith and Shackley, 2006). For a decade, this site has been the focus of native oyster restoration.

The Milford Haven Waterway (MHW) is a ria-estuary, an uncommon estuary type restricted in the UK to SW England, and Wales. The MHW is the only example of its kind in Wales and the largest ria-estuary complex in the UK (∼ 170 km long). Of the 55 km^2^ area covered, over 30% is composed of intertidal habitat (Burton, 2008). The Pembroke Marine Special Area of Conservation (SAC) is the UKs third largest SAC and includes almost the entirety of the Milford Haven Waterway. The native oyster *Ostrea edulis*, is part of the estuary feature for the SAC, and is a priority Biodiversity Action Plan species at a UK level. Milford Haven is host to one of only three surviving oyster beds in the U.K.

### Laboratory regime

Biometric data – wet weight (g), shell length (mm) and width (mm), position within a limpet stack – were recorded for each specimen. Sex was determined and assigned as either male (presence of developed penis), female (absence of penis), or transitionary (reduced penis) as described by Coe (1936). Haemolymph was collected from each limpet by carefully removing the flesh from the shell using a bunt-ended probe, allowing the haemolymph to pool in the shell cavity (**Fig. 1b**). Haemolymph was aspirated using a needle (23G) fitted onto a 1 mL sterile syringe. Total haemocyte counts (immune cells) were recorded using an improved Neubauer haemocytometer and optical microscope. Bacterial colony forming units (CFUs) were determined by spreading 100 uL of haemolymph diluted 1:100 with 3% NaCl solution on tryptone soya agar (TSA) plates supplemented with 2% NaCl (two (plates) technical replicates were performed per biological sample), and Thiosulfate-Citrate-Bile salts-Sucrose agar (TCBS). Plates were incubated at 25 °C for 24 h prior to counting CFUs. Bacterial loads are expressed as CFUs per mL haemolymph.

### Tissue histology

Whole tissue histology was used to screen a subset (n = 343) of animals to visualise any potential immune responses to vibrio-like bacteria, e.g., haemocyte infiltration or tissue damage. Whole tissue histology of *C. fornicata* was performed as described by Quinn et al. (2020). Briefly, samples were submerged in Davidson’s seawater fixative for 24h, rinsed in water and stored in 70% ethanol prior to processing. Samples were dehydrated using an ethanol series (70%-100%), transferred to Histoclear/Histochoice for 1 h, and infiltrated with molten wax using a Shandon™ automated tissue processor (Thermo Fisher Scientific, Altrincham, UK) prior to embedding. Sections were cut at 5–7 µm using a rotary microtome, adhered to glass slides using egg albumin, then stained with Cole’s haematoxylin and eosin. Slides were inspected using an Olympus BX41 photomicroscope. Images were adjusted for colour balance and contrast/brightness only.

### DNA extraction from limpets

A range of commercially available DNA extraction kits, namely the Sigma Aldrich GenElute ™ Blood Genomic DNA Kit, Qiagen DNeasy Blood and Tissue Kit, Qiagen DNeasy Powersoil Kit, the Omega Bio-TEK, E.Z.N.A® Tissue Kit, along with Chelex extraction, were trialled in order to extract DNA from the haemolymph of *C. fornicata*. DNA yields were quantified using the Invitrogen Qubit Fluorometer in combination with the Invitrogen dsDNA High Sensitivity Assay Kit. The Sigma Aldrich GenElute ™ Blood Genomic DNA Kit was selected for use throughout the sampling period based on DNA yields and quality. Genomic DNA from *C. fornicata* haemolymph (100 μL) was extracted following the supplier’s guidelines. For 29 out of 1800 specimens it was not possible to obtain 100 μL of haemolymph due to their small size, instead, 20 mg of solid foot tissue (largely muscle) was used.

### PCR-based assay for *Vibrio* spp

All PCR assays were carried out in 25 μL total reaction volumes using 2x Mastermix (New England Biolabs Inc., Ipswich, USA), 1.25 μL oligonucleotide primes (10 μM) synthesized by Eurofins (Ebersberg, Germany), 2 μL of genomic DNA, and performed in a PCR thermocycler (BioRad Laboratories Inc., Hemel Hempstead, UK.). For amplification of vibrios, the Vib1-F (GGCGTAAAGCGCATGCAGG) and Vib2-R (GAAATTCTACCCCCCTCTACAG) universal primer set (Vezzulli et al., 2012) was used and followed conditions described originally by Thompson et al. (2004) and modified by Vezzulli et al. (2012). The expected amplicon size was 113 bp. Negative controls consisted of DEPC-treated Molecular Biology Grade Water (Sigma Aldrich) in the absence of DNA template to avoid false positives due to contamination. Positive controls consisted of 1 μL DNA purified from the haemolymph of an infected donor crab (*Carcinus maenas;* Davies et al., 2021). PCR products were visualised using 2 % (w/v) agarose/TBE gels stained with 3 μL Greensafe premium nucleic acid stain (NZYTech, Lisboa, Portugal). TBE gels consisted of 100 mL 1x TBE buffer, and 2 g agarose. Each gel was run at 100 volts for 45 minutes in TBE buffer.

In preparation for sequencing positive signals, amplicons were purified using HT ExoSAP-IT™ Fast high-throughput PCR product clean-up (Thermo Fisher Scientific, Altrincham, UK). Direct Sanger sequencing was carried out by Eurofins. In total, 72 representative samples were chosen – distributed across both sites and from each month sampled. Of the 72 samples sent for sequencing, 22 of these returned usable data. Chromatograms were manually checked for mis-called bases to ensure the accuracy of the nucleotides. The generated sequences were trimmed manually of primer regions and matched against the National Center for Biotechnology Information (NCBI) nucleotide database using BLASTn (Basic Local Alignment Search Tool). Query Coverage (QC), Maximum Identity (MI) and E-value data were recorded for the top three returns per sample. Sequences were submitted to the NCBI short read archive (SRA) under accession numbers SRR13165025 – SRR13165046.

### Phylogenetic analysis

Retrieved sequences were searched against GenBank using default BLASTn settings followed by restricting the search to the genus *Vibrio*. Bacterial nucleotides derived from *C. fornicata* were added to a selection of known, geographically distributed vibrio reference sequences to make up a comprehensive dataset. A complete sequence alignment was achieved using the Clustal tool in MEGA X. Evolutionary analyses and reconstructions were carried out in MEGA X (Kumar et al. 2018) using the maximum likelihood routine based on the Kimura 2-parameter model and an independent Neighbour Joining routine. A consensus tree with the highest log likelihood value (–150.41) from 1,000 bootstrap re-samplings was formatted using iTOL (Letunic and Bork, 2019).

### Statistical analyses

To determine if specific predictor variables had any significant effect on the probability of finding vibrio*-*positive limpets, binomial logistic regression models were carried out using the Logit link functions found in the MASS library (following Bernoulli distributions). Logistic models were carried out in RStudio v1.2.5033 and R v4.0.2. An information theoretical approach was utilised for both model selection and assessment of model performance (Richards, 2005). Initial models will from herein be referred to as full models. Using the drop1 function, each non-significant predictor variable from the full model was removed sequentially to enhance the predictive power of the final model. The function of drop1 is to compare the initial full model with the same model, with the least significant predictor variable removed. If the difference between the reduced model and initial full model is different, the removed predictor variable is kept out of the new, reduced model. A Chi-square test is used for comparison of the residual sum of squares in both models in the case of binomial response variables. Variables included in the full model: vibrio (PCR negative/positive, 0 or 1), location (Milford Haven or Swansea Bay), season (Spring (Mar, Apr, May), Summer (Jun, Jul, Aug), Autumn (Sep, Oct, Nov), Winter (Nov, Dec, Jan), sex (male, female, transitionary), wet weight (continuous number), and position in stack (e.g., 1^st^, 2^nd^, 3^rd^, etc.). All other statistics (tests of normality, ANOVA, Kruskal-Wallis, Mann-Whitney, Chi-square) and figure preparation were performed in GraphPad Prism v.7/8 (GraphPad Software, La Jolla California USA).

## Results

### Population and biometric data of *Crepidula fornicata*

Across the three morphometric measures, i.e., wet weight (g), shell length (mm) and width (mm), limpets dredged from Swansea Bay were significantly larger (Fig. 2; P < 0.0001, Mann-Whitney). For weight, Milford Haven *C. fornicata* were 7.99 ± 0.10 g (range, 0.4 to 20.17 g) – which is less than half the mass of those from Swansea Bay, 18.86 ± 0.3 g (range, 0.29 to 70.1 g; Fig. 2a). Similar observations were made with respect to length (Fig. 2b) and width (Fig. 2c): *C. fornicata* from Milford Haven were 36.1 ± 0.2 mm (range, 10 to 53 mm) and 23.9 ± 0.1 mm (range, 11 to 43 mm), respectively, and 46.5 ± 0.3 mm (range,14 to 61 mm) and 27.8 ± 0.1 mm (range, 11 to 42 mm) from Swansea Bay, respectively.

**Fig. 2.**
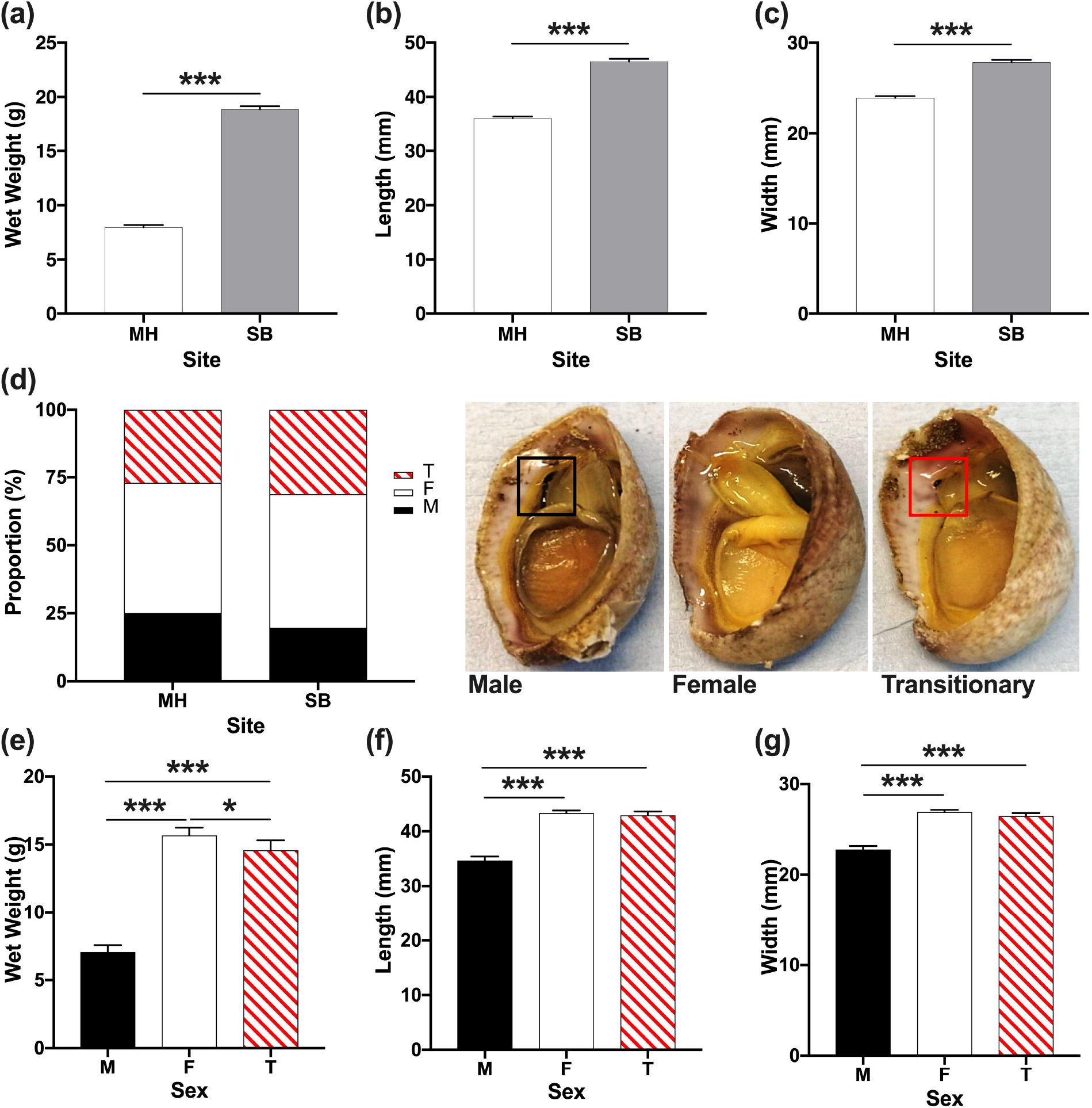
Morphometric measurements of *C. fornicata*. Overall wet weight **(a)**, length **(b)** and width **(c)** of limpets according to site, Milford Haven (MH, *n* = 900) and Swansea Bay (SB, *n* = 900). Proportions of male (M), female (F) and transitionary (F) limpets according to site **(d)**. Inset, the presence of a fully formed penis (black box) indicates the individual is male, the absence of a penis indicates a female, and the presence of a reduced penis (red box) indicates a transitionary state. Average weights **(e)**, lengths **(f)** and widths (g) of limpets according to sex. Asterisk(s) denotes significant differences (*, P < 0.01; ***, P < 0.0001). Data are expressed as mean values with 95% CI, *n* = 1,800.

At both sites, similar proportions of males, females, and transitionary individuals made-up each population. Females dominated significantly more so than either males or transitionary limpets (Fig. 2d; *Χ*^2^(2, n=1800) = 8.852, *P* = 0.012; Chi-square). At Milford Haven, females accounted for 48%, males 25%, and transitionary 27%. At Swansea Bay, females made up 49%, males 20%, and transitionary 31%. Not only were there fewer males in general, but female and transitionary *C. fornicata* were found to be significantly heavier (*P* < 0.01, Kruskal-Wallis; Fig. 2e) - with females also heavier than transitionary individuals (*P* = 0.013). Females weighed 15.67 ± 0.29 g (range, 2.69 to 70.10 g), followed by transitionary at 14.58 ± 0.37 g (range, 0.57 to 39.61 g), then males 7.06 ± 0.26 g (range, 0.29 to 41.34 g). This trend was also reflected in the average shell lengths (Fig. 2f) and widths (Fig. 2g) among the sexes – females and transitionary limpets were significantly longer and wider when compared to males (*P <* 0.001, Kruskal-Wallis; Fig. 2).

*Crepidula fornicata* are known to form stacks on hard surfaces or basibionts (e.g., oyster; Figs. 3a and 3b). Stack sizes of limpets from Swansea Bay (1-11) were consistently, and significantly (*Χ*^2^(10, N = 1800) = 95.0, *P* < 0.0001; Chi-square test), larger than those found in Milford Haven (1-5). At the latter site, 50% of stacks consisted of just one individual, 35% consisted of 2 individuals, 12% consisted of 3 individuals, and the remaining 3% was distributed between stacks of 4 and 5. At Swansea Bay, 40% of stacks consisted of one individual, followed by 25% of 2, 13% of 3, 3.8% of 4, and the remaining 14% was distributed among stacks of 5 to 11 limpets. A clear temporal trend was observed regarding the number of *C. fornicata* that formed stacks (*P* < 0.0001; Kruskal-Wallis) with the smallest encountered during the summer, ranging from 1 to 7 limpets (n = 450), and largest during the winter, ranging from 1 to 11 (n = 440). Generally, weight, length and width of limpets decreased the further up the stack (Figs. 3c, 3d). Within stack position was found to influence the wet weight of an individual, with those at the bottom being significantly larger in size than those above (*P* < 0.0001, Kruskal-Wallis).

**Fig. 3.**
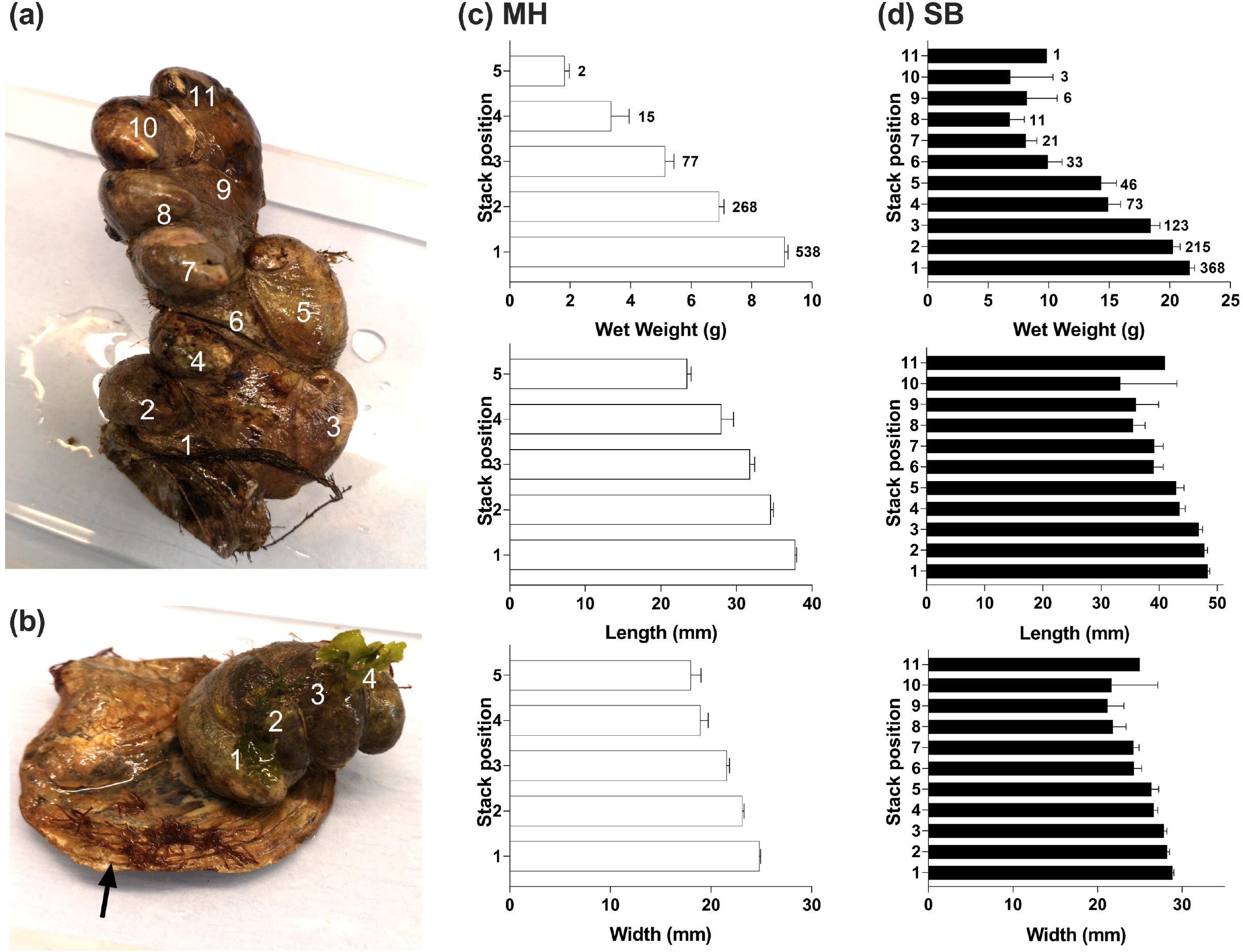
Differential stack formation of *C. fornicata* across two sites in South Wales, UK. **(a)** *In extremis*, limpets can form stacks up to 11 individuals – these limpets were photographed from Swansea Bay, January 2019. **(b)** Routinely, limpets from stacks on hard substrates including mussel and oyster (*Ostrea edulis*, black arrow) shells, i.e., basibionts. Morphometric data (weight, length, width) categorised according to position within stacks for Milford Haven **(c)** and Swansea Bay **(d)**. In the two upper (middle, right) panels for weight, sample size frequencies are presented for each position within the stack (n = 900 per site, 1,800 in total).

Freely circulating, homogeneous immune cell (haemocyte) populations were present in the haemolymph of *C. fornicata* and quantified from a total of 1,771 individuals (Milford Haven *n* = 883, Swansea Bay *n* = 888; Fig. 4a-c). Overall, significantly fewer haemocytes were present in those limpets from Milford Haven, 2.7 ×10^6^ ± 9.3 ×10^4^ mL^-1^ (1.3 ×10^5^ to 3.3 ×10^7^ mL^-1^), when compared to Swansea Bay, 3.3 ×10^6^ ± 1.1 ×10^5^ mL^-1^ (3 ×10^5^ to 3.1 ×10^7^ mL^-1^; *P* < 0.01; Fig. 4a). Temporal fluctuations in haemocyte numbers were apparent, with season (F_(3, 1761)_ = 21.56, P < 0.001) and site (F_(1, 1761)_ = 17.96, P < 0.001) representing significant contributing factors. Haemocyte levels peaked in the summer/autumn, ranging from ∼2.8 ×10^5^ to 3.3 ×10^7^ mL^-1^, and were at their nadir in spring for Swansea Bay and winter for Milford Haven, ranging from ∼1.3 ×10^5^ to 1.6 ×10^7^ HC/mL (Fig. 4b). When combining both sites, limpets collected in March recorded the lowest haemocyte counts of 1.79 ×10^6^ ± 1.75 ×10^5^ mL^-1^ (from 1.3 ×10^5^ to 1.47 ×10^7^ mL^-1^), whereas haemocyte numbers peaked in September at 4.64 ×10^6^ ± 3.91 ×10^5^ mL^-1^ (from 5.1 ×10^5^ to 3.1 ×10^7^ mL^-1^). Generally, haemograms increased in limpets positioned further up the stack (**Supplementary Figs. 1a and 1b**). This trend was only observed between individuals in position 1 and position 6 (P = 0.0453) when data for each site were combined.

**Fig. 4.**
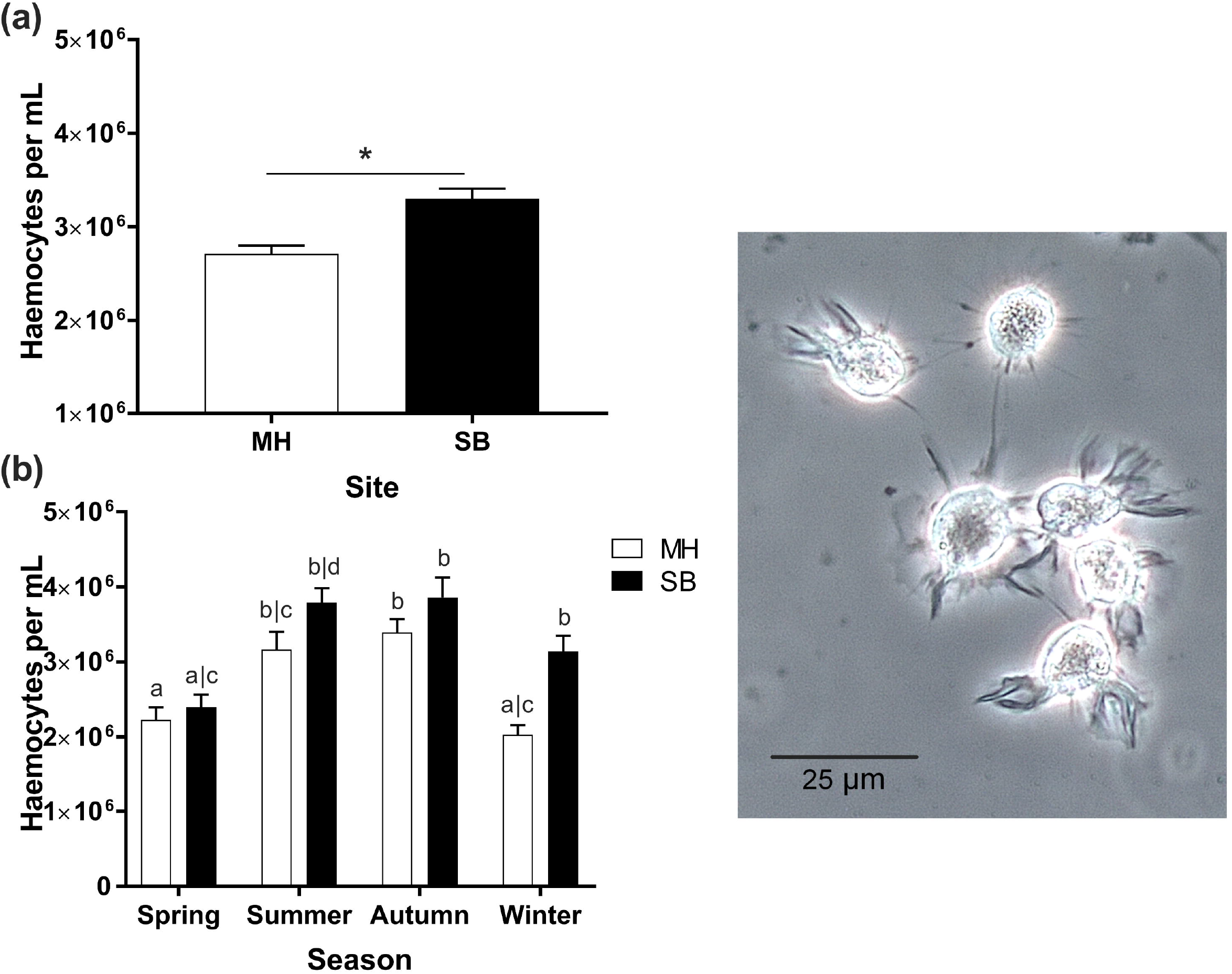
Total haemocyte counts observed in the haemolymph of *C. fornicata*. **(a)** Average numbers of haemocytes per mL in limpets collected from Milford Haven (MH, = 883) and Swansea Bay (SB, n = 888). **(b)** Temporal profiles of haemocyte numbers according to site and season. Unshared letters denote significant differences (P < 0.05), and the asterisk (*) represents P < 0.01. Inset, phase contrast image depicting the typical appearance of haemocytes (granular, motile, refractile).

### Spatiotemporal dynamics of vibrio-like bacteria hosted by *Crepidula fornicate*

Sufficient volume of haemolymph was retrieved from 1,780 individuals and plated on the generalist growth medium TSA (+ 2% NaCl), and on the vibrio selective TCBS (**Fig. 5**). Of these samples, <1% of limpets contained no signs of cultivable bacterial colonies on either TSA (n = 8) or TCBS (n = 12). When combining data from both sites, average bacterial growth on TSA was 2.1 ×10^5^ ± 7.1 ×10^3^ CFUs/mL (range = 0 to 3.8 ×10^6^ CFU/mL), while on TCBS numbers reached 1.8 ×10^5^ ± 6.6 ×10^3^ CFUs/ mL (range = 0 to 2.69 ×10^6^ CFU/mL). No significant differences in bacterial loads from the haemolymph of *C. fornicata* were detected between Milford Haven and Swansea Bay for either TSA (*P* = 0.487, Mann-Whitney; **Fig. 5a**) or TCBS (*P* = 0.665; **Fig. 5b**). A clear temporal trend was detected among haemolymph CFUs from each location across the 12-month survey (**Fig. 5**). When data were combined from both sites, culturable bacterial numbers were at their highest during the summer (**Figs. 5c and 5d**) **–** 4.1 ×10^5^ ± 1.76 ×10^4^ CFU/mL (range 3.5 ×10^3^ to 2.6 ×10^6^ CFU/mL) for TSA and 3.99 ×10^5^ ± 1.88×10^4^ CFU/mL (range 0 to 2.7 ×10^6^ CFU/mL) for TCBS. Bacterial presence was lowest during winter, again for both TSA and TCBS media, 1.12 ×10^5^ ± 7.7 ×10^3^ CFU/mL (range 0 to 1.1 ×10^6^ CFU/mL) and 7.3 ×10^4^ ± 6.74 ×10^3^ CFU/mL (range 5 ×10^2^ to 7 ×10^5^ CFU/mL), respectively. There were some subtle differences in seasonal CFU highs/lows when comparing each site and growth medium (**Figs. 5e** and **5f**). Limpets collected in August displayed the highest CFU numbers on TSA and TCBS (**Supplementary Figs. 2a and 2b**), ∼6 to 7 ×10^5^ CFU/mL (range 3 ×10^4^ to 2 ×10^6^ CFU/mL), whereas those collected during the winter months contained the fewest on TSA (7.29 ×10^4^ CFU/mL; range 4 ×10^3^ to 7 ×10^5^ CFU/mL) and TCBS (3.56 ×10^4^ ± 5.87 ×10^3^ CFU/mL; range 5 ×10^2^ to 4.48 ×10^5^ CFU/mL), respectively.

**Fig. 5.**
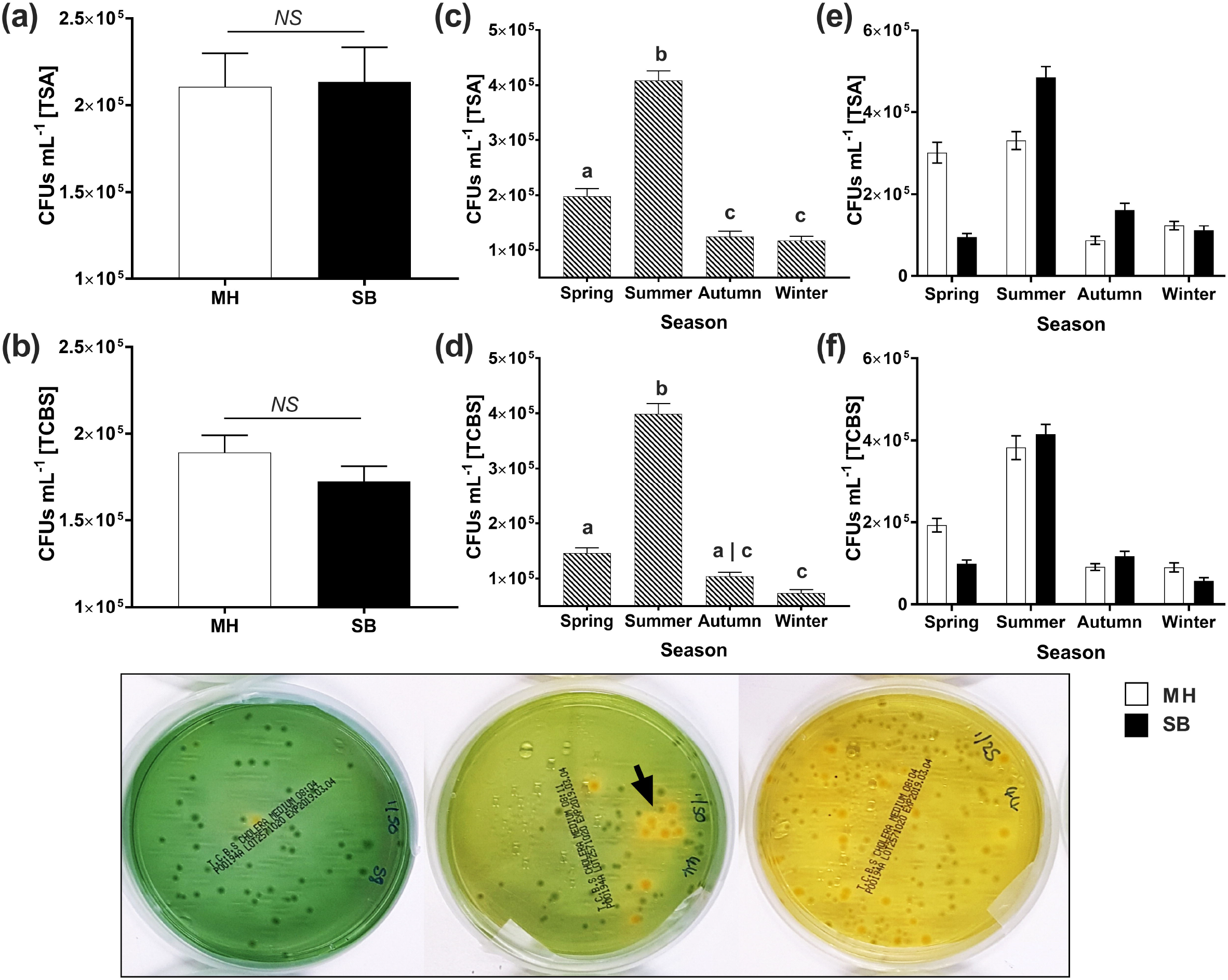
Spatiotemporal patterns of cultivable vibrio-like bacteria from the haemolymph of *C. fornicata*. Average number of bacterial CFUs per mL haemolymph on TSA **(a)** and TCBS **(b)**. Average number of bacterial CFUs across seasons on TSA **(c)** and TCBS **(d)**. Average number of bacterial CFUs, per site, per season on TSA **(e)** and TCBS **(f)**. Individuals were sampled from Swansea Bay (SB) and Milford Haven (MH). *C. fornicata* haemolymph was collected, diluted 1:100 with sterile 3% saline, streaked (100 µL) onto the respective agar plate, and incubated at 25 °C for 24 h prior to counting. NS, non-significant (P > 0.05) differences. Unshared letters represent significant differences (P < 0.05) determined from Tukey’s post-hoc tests (n = 1,780). **Inset**, representative growth of cultivable bacteria from haemolymph of *C. fornicata* incubated at 25 °C for 24 h on TCBS - *Vibrio* selective agar. The yellow colour change is an indication of pH change due to the fermentation of sucrose (black arrow). Bacteria not capable of fermenting sucrose produce green to blue-green colonies.

We further investigated whether the location of a limpet within a stack or the number of haemocytes in the haemolymph were linked to bacterial CFU counts. Stack position of a limpet was a non-significant contributing factor to CFU counts on TSA (*P* = 0.32) or TCBS (*P* = 0.063; **Supplementary Figs. 3a and 3b**). Overall, a positive relationship was observed between the bacterial burden (CFUs mL^-1^) of *C. fornicata* haemolymph and the number of haemocytes from the corresponding limpet (**Supplementary Figs. 4a and 4b**) *–* i.e., increases in CFUs and haemocyte numbers were reciprocal. When data were analysed per site, the positive correlation was non-significant on either media (TSA, *P* = 0.978; TCBS, *P* = 0.157) for the limpets from Milford Haven (Fig. 6a). Conversely for the limpets from Swansea Bay, the positive correlation between haemolymph CFUs and haemocytes was significant (P < 0.001, Spearman correlation; **Fig. 6b**).

**Fig. 6.**
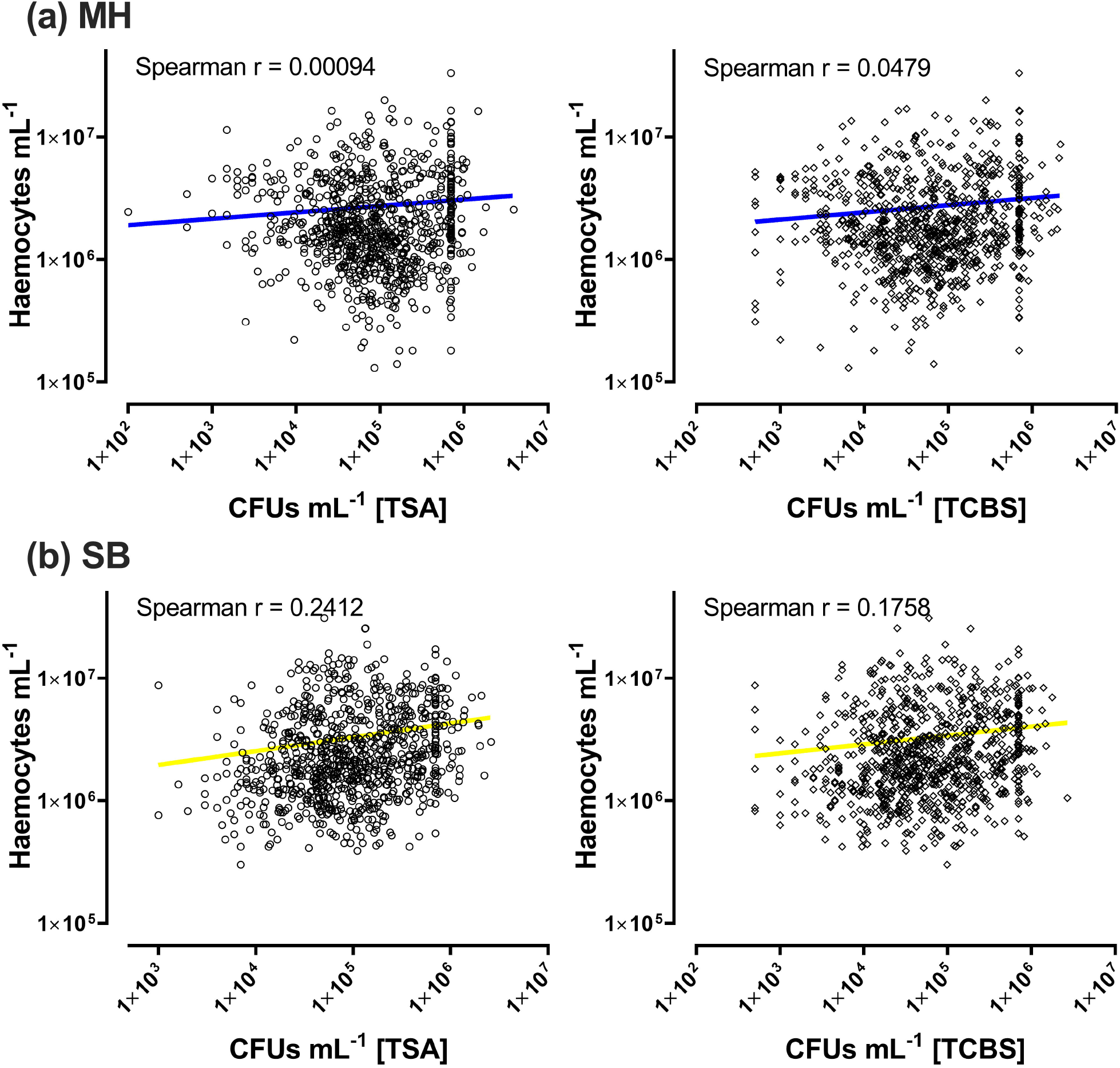
Relationship between bacterial colony forming units (CFU/mL) and haemocytes in the haemolymph of *C. fornicata*. (a) *C. fornicata* collected from Milford Haven (MH; n = 883) - haemolymph plated on generalist TSA (upper left panel) and vibrio-selective TCBS (upper right panel). **(b)** *C. fornicata* collected from Swansea Bay (SB; n = 887) - haemolymph plated on generalist TSA (lower left panel) and vibrio-selective TCBS (lower right panel). Bacterial load is expressed as CFUs per mL haemolymph.

All 1,800 limpets were screened for the presence of *Vibrio* spp. using end-point PCR, with 83% testing positive overall (single amplicons of the expected size on agarose gel electrophoresis; **Supplementary Fig. 5**). When considering site, detection rates were 85% (763/900) and 81% (729/900) for Swansea Bay and Milford Haven, respectively (**Fig. 7a**). August was the month with the highest number of PCR-positive samples at 150/150 (100%), whereas the fewest PCR-positive limpets were in February with 84/150 (56%; **Fig.7b**). Biometric data recorded during the processing of *C. fornicata* (size, sex, position in stack), along with collection site, bacterial CFU counts, and haemocyte levels were incorporated into Binomial Logistic Regression models to determine whether any of these factors contributed to the occurrence/detection of *Vibrio* spp. *via* PCR. Limpet wet weight (**Fig.7c**), season (**Fig. 7d**) and TCBS counts (**Fig. 7e**) are significant predictor variables (**Table 1**). Significant decreases in the number of vibrio*-*positive limpets were observed during Spring (*P =* 0.00539) and Winter (*P =* 0.0064). Individuals with more bacterial colonies on selective TCBS medium were most likely to test positive through for *Vibrio* spp. than those individuals that were PCR-negative (*P <* 0.0001; ∼2 ×10^5^ versus ∼9 ×10^4^ CFU/mL, respectively).

**Table 1.** Binomial logistic regression models testing the influence of biometric and environmental predictor variables on the presence of vibrio-like bacteria in *C. fornicata* sampled from Swansea Bay and Milford Haven (Jan 19 – Dec 2019).

| Model | Predictor variable | Estimate slope | SE | P-value |
| --- | --- | --- | --- | --- |
| **Full Model**<br>Vibrio ~ Site + Season + Sex + Position.In.Stack + Wet.Weight + TSA + TCBS + Haemocyte.Counts | SiteSB | 0.03827 | 0.1913 | 0.8414 |
|  | SeasonSpring | -0.5164 | 0.1878 | **0.0059\*\*** |
|  | SeasonSummer | -0.07122 | 0.2272 | 0.7539 |
|  | SeasonWinter | -0.5723 | 0.1894 | **0.0025\*\*** |
|  | SexM | -0.275 | 0.1955 | 0.1596 |
|  | SexT | -0.0008116 | 0.1575 | 0.9958 |
|  | Position.In.Stack | 0.05505 | 0.05704 | 0.3345 |
|  | Wet.Weight | 0.02108 | 0.01263 | 0.0949 |
|  | TSA | 4.455E-07 | 4.609E-07 | 0.3338 |
|  | TCBS | 0.000001913 | 5.835E-07 | **0.0010\*\*** |
|  | Haemocyte.Counts | 1.53E-10 | 2.329E-08 | 0.9947 |
| **Model reduced 4** |  |  |  |  |
| **Vibrio ~ Season + Wet.Weight + TCBS** | SeasonSpring | -5.06E-01 | 1.82E-01 | **0.0053\*\*** |
|  | SeasonSummer | -4.87E-02 | 2.23E-01 | 0.8269 |
|  | SeasonWinter | -4.92E-01 | 1.80E-01 | **0.0064\*\*** |
|  | Wet.Weight | 2.85E-02 | 8.70E-03 | **0.0010\*\*** |
|  | TCBS | 2.24E-06 | 4.45E-07 | **4.97E-07\*\*\*** |

Significance. codes: 0 '\*\*\*' 0.001 '\*\*' 0.01 '\*' 0.05 '.' 0.1 ' ' 1 #Vibrio = PCR positive

**Fig. 7.**
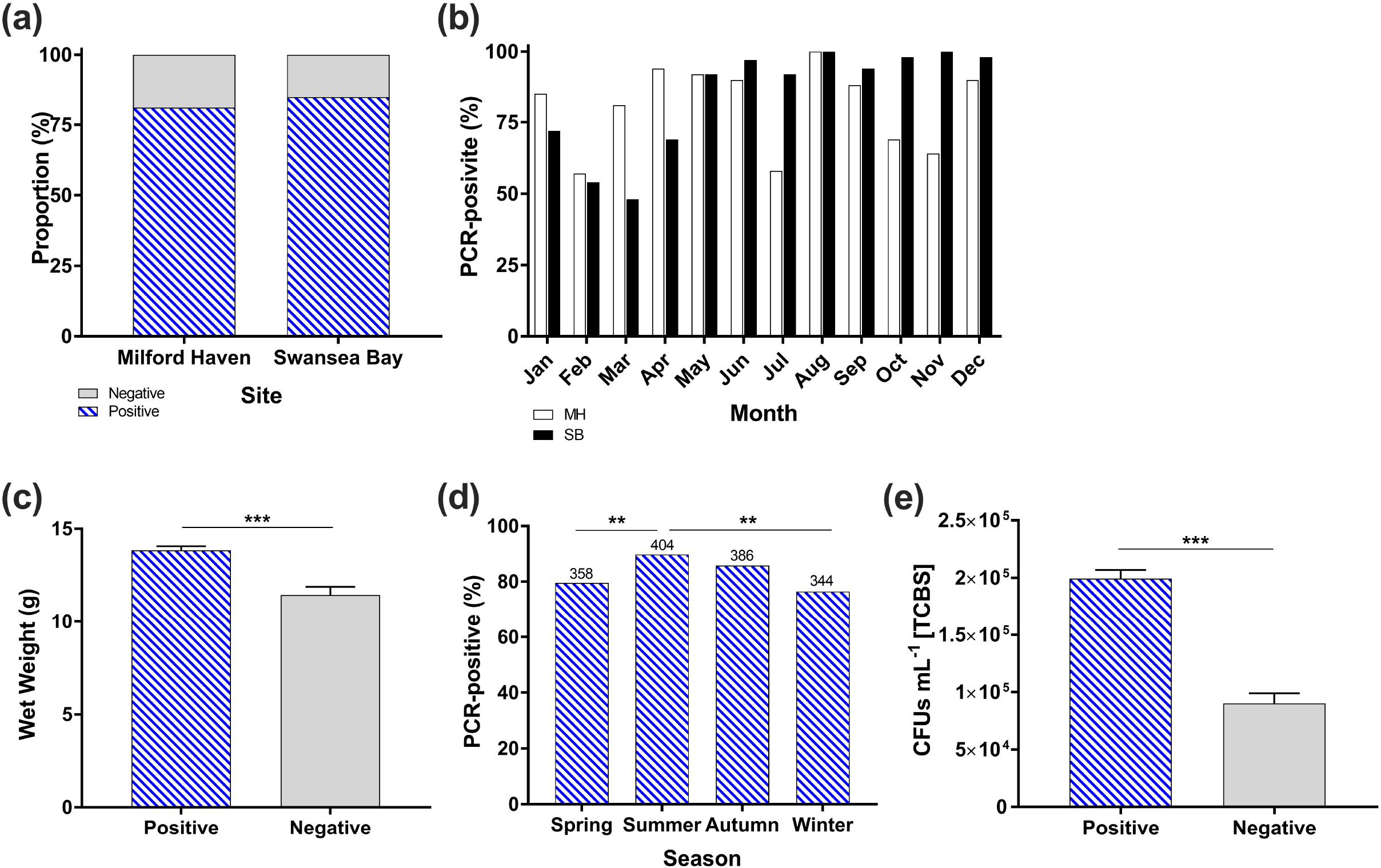
Prevalence of vibrio-like bacteria among *C. fornicata* determined *via* PCR-based screening. Limpets detected positive for *Vibrio* spp. across sites **(a)** and months **(b)**. Haemolymph samples were used for 1771/1800 individuals, with solid tissue samples being used for 29/1800 due to their small size. Of the 29 solid tissue samples, 14 yielded PCR positive bands for vibrio-like bacteria, with 12 of these limpets containing cultivable bacteria from their haemolymph (grown on TSA /TCBS). Panels **(c), (d)** and **(e)** represent the significant predictor variables determined using Binomial Logistic Regression Models (see Table 1), limpet weight, season and CFU counts on vibrio selective TCBS, respectively. Significant differences are denoted by asterisks (** P < 0.01, ***, P < 0.001). In panel (d), the number of positive individuals from each respective season are placed above each column.

### Identification of vibrio-like bacteria

A selection of vibrio-positive amplicons (n = 72) from both sampling sites and each month were sent for direct sequencing using Sanger’s method in a bid to identify putative vibrios to species level (although the target region is small, 113 bp). Of these 72 samples, 22 high quality sequences were generated with 7 proving clear matches to known *Vibrio* species, with the remainder matching unclassified/uncultured vibrio-like bacteria (**Supplementary Table 1**). BLASTn searches revealed consistent matches (85-100% coverage/identity) with members of the Splendidus clade: *V. tasmaniensis, V. celticus, V. crassostreae, V. gallaecicus, V. kanaloae*, and *V. toranzoniae*. Alignments and phylogenetic trees of the partial 16S ribosomal RNA sequences displayed consistent topology when reconstructed using maximum likelihood and neighbour-joining routines – indicating the presence of some unique vibrio sequence types in *C. fornicata* (**Fig. 8**). Trees were comprised of vibrios isolated from diseased and healthy molluscs, as well as environmental isolates. A distinct Splendidus clade was formed with key representative vibrios, as well as sequences amplified from *C. fornicata*. A second, distinct well supported (99%) grouping of vibrio-like sequences from both Milford Haven and Swansea Bay sites clustered together, thereby suggesting at least two vibrio ecotypes are present in these limpets. Sequences were distributed among benign and pathogenic bacteria. Due to the small size of the final reads (∼60 bp), caution is needed when interpreting the potential species identity of these bacteria, however, they do represent members of the *Vibrio* genus and Splendidus clade.

**Fig. 8.**
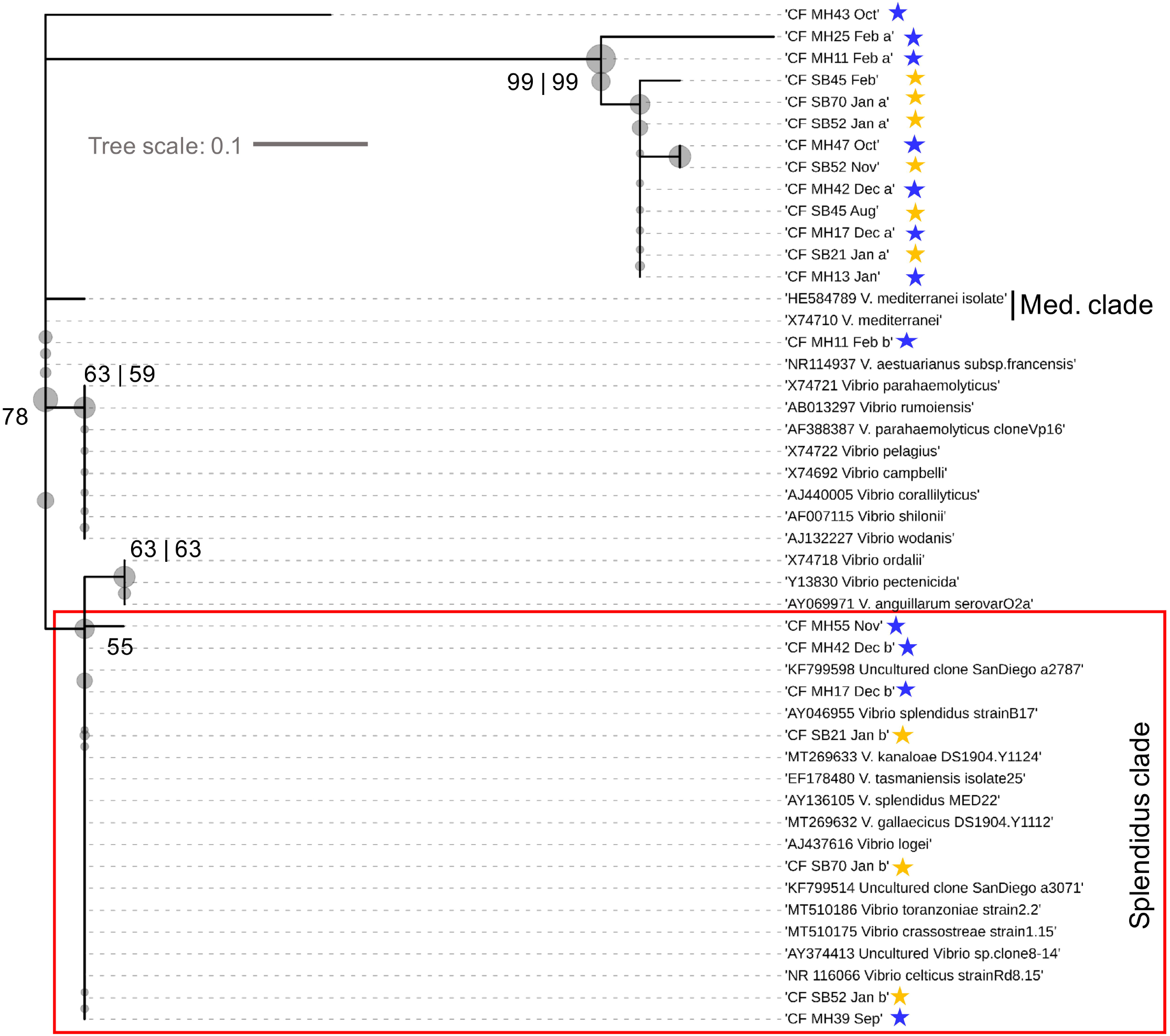
Consensus phylogram (unrooted) of the partial 16S rRNA region recovered from slipper limpets *Crepidula fornicata* (SRR13165025 – SRR13165046). Reference sequences represent the top BLASTn-search results from NCBI plus a broad list of known, geographically distributed *Vibrio* species. Bootstrap support values are depicted as spheres (ML | NJ). The scale represents nucleotide substitutions per site (maximum likelihood (ML) estimation, 1000 bootstrap replicates). All reference sequences used were ≥90% (coverage and identity) matches for the sequences retrieved from limpets. Stars represent sites (blue, Milford Haven; yellow, Swansea Bay). The red box highlights a putative vibrio ecotype within the Splendidus clade.

### Bacteria-limpet interactions

A minimum of 20 individuals per month across both sites (except Jan, n = 19) were processed for multi-tissue histology (n = 343 in total, including those limpets with high bacterial loads or with no signs of bacterial presence). Ostensibly, there was no gross evidence of bacterial infection, tissue-wide damage and necrosis, or inflammation in the slipper limpets examined, regardless of their haemolymph bacterial load. In very few individuals, there were discrete signs of immune cell (haemocyte) infiltration, nodule formation, and varied states of what appeared to be compromised tissue – however, these could not be attributed to microbial disease (bacteriosis). There was one exception – a limpet from Milford Haven (#40 May 2019) that was undergoing regional bacterial putrefaction in some of the tissues (**Fig. 9**). Affected tissues appeared necrotic and likely became infested with bacteria consequently. Variable morphologies of bacteria were apparent (e.g., heterogeneous rod-shaped) in connective tissue (**Fig.9a**), which represents a clear sign that this is not a bacterial disease, i.e., vibriosis, but early putrefaction of a moribund animal. The digestive gland was replete with cellular fragments/debris reminiscent of epithelial sloughing and cell death (**Fig. 9b**). In healthy limpets, gill filaments formed a continuous, highly ciliated layer with discrete luminal spaces (**Fig. 9c**). The gills from the septic/moribund animal showed signs of damage to the outer epithelium and many cells in the space around the gills (e.g., lacking cilia), perhaps representing sloughed cells and/or tissue disintegration (**Fig. 9d**). Lipofuscin-containing granule abundance and intensity can vary greatly across tissues within an individual, and so it is unclear whether this represents a pathological condition/infection (some links to metal detoxification). The septic limpet depicted heterogeneous accumulation of lipofuscin associated with compromised tissues, e.g., distal gill filaments (**Fig. 9e**).

**Fig. 9.**
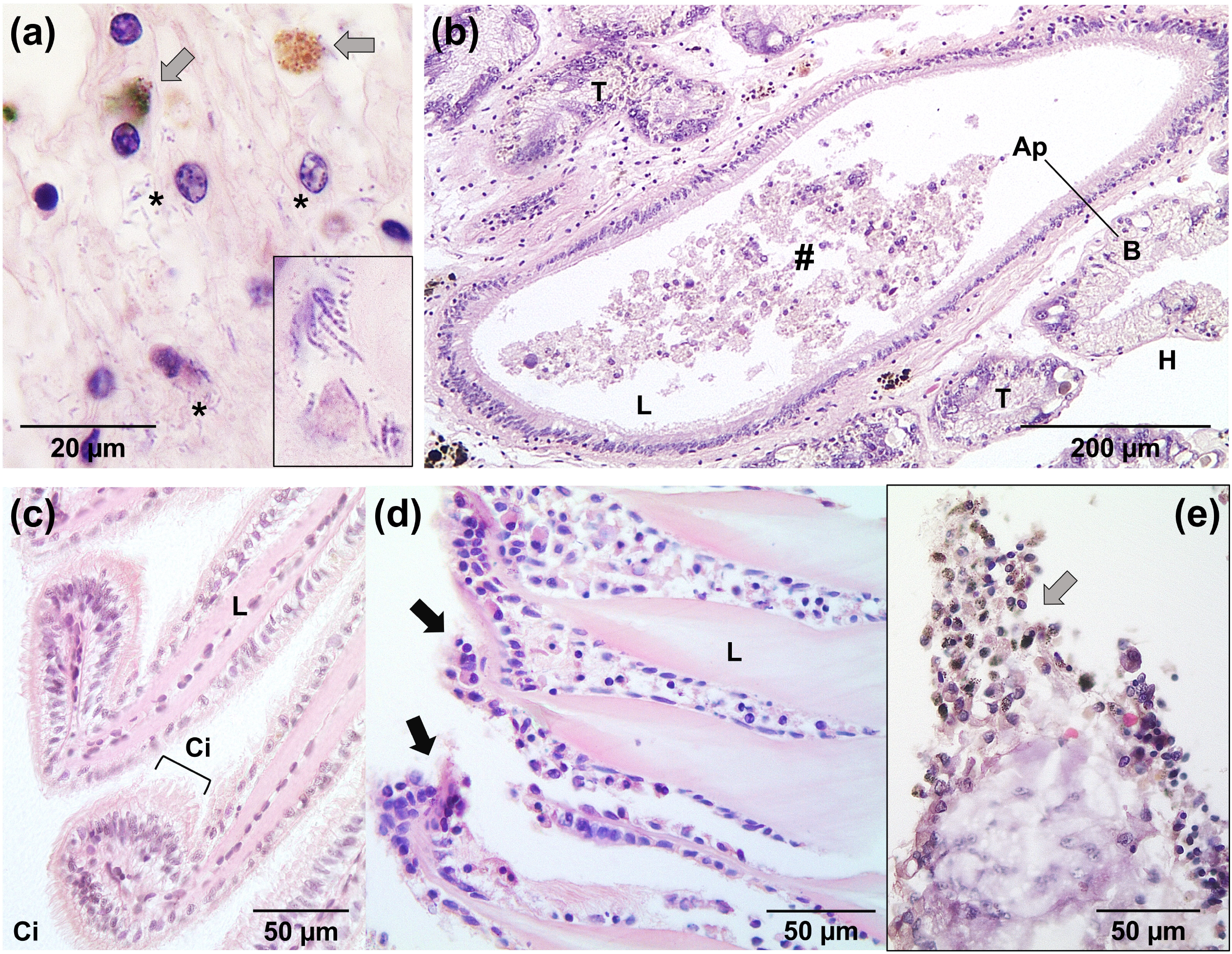
Histopathology of a moribund slipper limpet. **(a)** Septic connective tissue with various rod-shaped bacterial consortia (indicated by asterisks, *). Lipofuscin granules are denoted by grey arrows. Inset, magnified region of connective tissue replete with bacteria. **(b)** View of the intact digestive gland depicting gross debris accumulation (#) within the lumen (L). Ap, apical; B, basolateral; H, haemocoel (body cavity); T, tubule. **(c)** Healthy gill tissue with intact cilia (Ci) and regular lumen (L) architecture. **(d)** Compromised gill tissue lacking cilia (black arrows) (L). **(e)** Degenerating distal gill filament with gross accumulation of lipofuscin-like granules with melanin deposits (grey arrow) and some necrosis. All panels represent limpet #40 collected from Milford Haven in June 2019, except panel (c), which is from a seemingly healthy limpet from the same month.

## Discussion

To date, little to no disease or parasites have been reported in *C. fornicata* other than the presence of the shell boring sponge *C. celata* (Le Cam and Viard, 2011) and interactions with trematodes (Pechenik et al., 2001; Thieltges et al., 2006 and 2008). Herein we determined the bacterial loads of the invasive slipper limpet *C. fornicata*, focusing on a key aetiological agent in commercial shellfish, namely vibrios. Slipper limpets from two survey sites contained high levels, 83 to >99%, of vibrios in the haemolymph, but these bacteria were not pathogenic to *C. fornicata* (determined via histology). Instead, slipper limpets seemed to accumulate vibrio-like bacteria passively due to their filter feeding nature, thereby acting as a sink for these autochthonous microbes. Such observations have been made previously for marine invertebrates (Froelich and Oliver, 2013). Bivalves, such as clams, can be found with levels of vibrios in their tissue of 10^5^ CFU/g or higher, this being up to 100-fold greater than the surrounding waters (Froelich et al., 2017). Larger limpets were statistically more likely to contain vibrios, and at higher levels. This could be a consequence of age – positioned toward the bottom of a stack where younger/smaller limpets pile on top and remain *in situ* for extended periods of time.

Not all members of the *Vibrionaceae* are a threat to shellfish or humans – the genus *Vibrio* constitutes free-living bacteria, opportunists, pathobionts, and symbionts (e.g., McFall-Ngai, 2014; Wendling et al., 2014; Destoumieux-Garzón et al., 2020). A great diversity of *Vibrio* species, strains and ecotypes can be found among the microflora of marine invertebrates, e.g., spider crabs *Maja brachydactyla* (Gomez-Gil et al., 2010). Much work has been performed on the environmental drivers of dysbiosis in commercial shellfish, like the Pacific Oyster (*Crassostrea gigas*), linked to vibriosis and mass mortality events that can devastate populations (Destoumieux-Garzón et al., 2020). Despite some morphometrical differences observed between *C. fornicata* collected at Swansea Bay and Milford Haven, ‘site (or location)’ was not a significant factor when it came to the probability of a limpet testing positive for vibrios. Instead, seasonality was a key contributing factor. The highest prevalence of vibrio-like bacteria occurred during summer and late autumn. This is not entirely surprising as higher temperatures, especially those exceeding 15-17°C, enable vibrios to thrive and are linked to disease outbreaks and the expression of virulence factors (Stauder et al., 2010; Vezullli et al., 2012 and 2013). In addition to season, both weight and the presence of CFU growth on TCBS medium, were key predictors of bacterial presence. Criticism has been levelled at the use of TCBS medium for monitoring vibrios beyond key pathogens, such as *V. cholerae* and *V. parahaemolyticus* (Cavallo and Stabili, 2002; Donovan and van Netten, 1995; Gyraite et al., 2019). This medium was originally developed to enumerate vibrios from human and environmental samples (Kobayashi et al. 1963), however, it is apparent that not all vibrios grow on this medium and that some other taxa like aeromonads can grow (Lotz et al. 1983). For example, in a study of bacteria isolated from samples from estuarine sites only 61% of colonies taken from TCBS plates were found to be vibrios (Pfeffer and Oliver 2003). Even considering the putative drawbacks of using TCBS agar for vibrio enumeration in haemolymph, our results found that CFU growth on TCBS aligned with PCR outputs, thereby representing a validated approach for monitoring vibrio-limpet epizootiology as the coastal waters become warmer (or limpets move further outside of their native range). The frequency of vibrio infections has been increasing worldwide, between 1996 and 2006, the average annual incidence rose by 78%, and between 2008 and 2018 the incidence rate increased by 272% (Froelich and Daines, 2020). In the summer of 2015, Canada experienced the highest number of reported cases of human *V. parahaemolyticus* infections ever reported from the consumption of raw oysters (Taylor et al. 2018). This outbreak was associated with higher-than-normal sea surface temperatures (SST > 15°C), and the incidence rates decreased once SST dropped below 15°C (Taylor et al. 2018).

Limpets gathered from both our survey sites are co-located with high value shellfish, e.g., native oyster (*Ostrea edulis*) in Swansea Bay, therefore, part of our study was to determine the potential connectivity of disease, i.e., whether slipper limpets represent an overlooked reservoir for vibrios. PCR-based confirmation of vibrio presence using the universal Vib1-F/Vib2-R primer set was associated significantly with TCBS-based vibrio detection (Table 1), however, the amplicons generated were rather short at ∼60-100 bp in length, making it difficult to identify specific vibrios down to species level. Nonetheless, high quality sequences were retrieved across sites and months and identified as members of the Splendidus clade (e.g., *V. tasmaniensis, V. celticus*), and putative members of the Mediterranean clade. The former clade is commonly encountered among the ‘vibriome’ of marine animals, e.g., >21–77% of bacterial isolates of spider crab haemolymph sampled along the coastal waters of north-west Spain and the Canary Islands (Gomez-Gil et al., 2010). We did not encounter evidence to suggest that major pathogens of humans (e.g., *V. cholerae*) or shellfish (e.g., *V. aestuarianus*) are enriched within the bacterial consortia of slipper limpet haemolymph. Follow-on efforts to characterise vibrios should use high throughout sequencing or long read amplification.

## Concluding remarks

Findings from our study go some way to address the knowledge deficit with respect to the disease profile of invasive *C. fornicata*, alongside the identification of key variables that influence the presence of vibrio in limpets across regions in South Wales, UK (age and seasonality). Strikingly, we encountered a single moribund animal out of 1800 screened with several tissues partially putrefied and replete with bacterial morphotypes, despite the prevalence and severity of vibrios in the limpet haemolymph. It is possible that the vibrios present in limpet haemolymph provide an element of protection through competitive exclusion of other disease-causing agents. We consider that slipper limpets are not highly susceptible to bacteriosis at either site surveyed, and do not harbour vibrios known to be pathogenic to humans. The lack of susceptibility to local pathogenic bacteria may explain, in part, the invasion success of *C. fornicata* across this region.

## Acknowledgements

We should like to thank skippers Mr Keith Naylor and Mr Max Robinson (R.V. Mary Anning, Swansea University) and Mr Barry Thomas for their assistance with sample collection in the Swansea Bay area.

## Declarations

### Author contributions

Conceptualization, CC. Data curation, EQ, SM, JT, AR, CC. Formal analysis, EQ, AR, CC. Funding acquisition, AR, CC. Investigation, EQ, JT, SM, RP, CD, AR, CC. Methodology, EQ, JT, SM, CD, AR, CC. Project administration, SM, AR, CC. Resources, AR, CC. Software, EQ, CD, AR, CC. Supervision, AR, CC. Validation, EQ, AR, CC. Visualisation, EQ, CC. Writing – original draft presentation, reviewing and editing, EQ, AR, CC.

### Availability of data and material

Sequence data has been deposited into NCBI’s short read archive (SRA) under accession numbers SRR13165025 – SRR13165046.

### Code availability

Not applicable.

### Ethics approval

Sampling and experimental work on *C. fornicata* were approved by the College of Science (Swansea University) research ethics committee, SU-Ethics-111217/446.

