## Supplementary information for "Invasive slipper limpets *Crepidula fornicata* act like a sink, rather than source, of *Vibrio* spp."

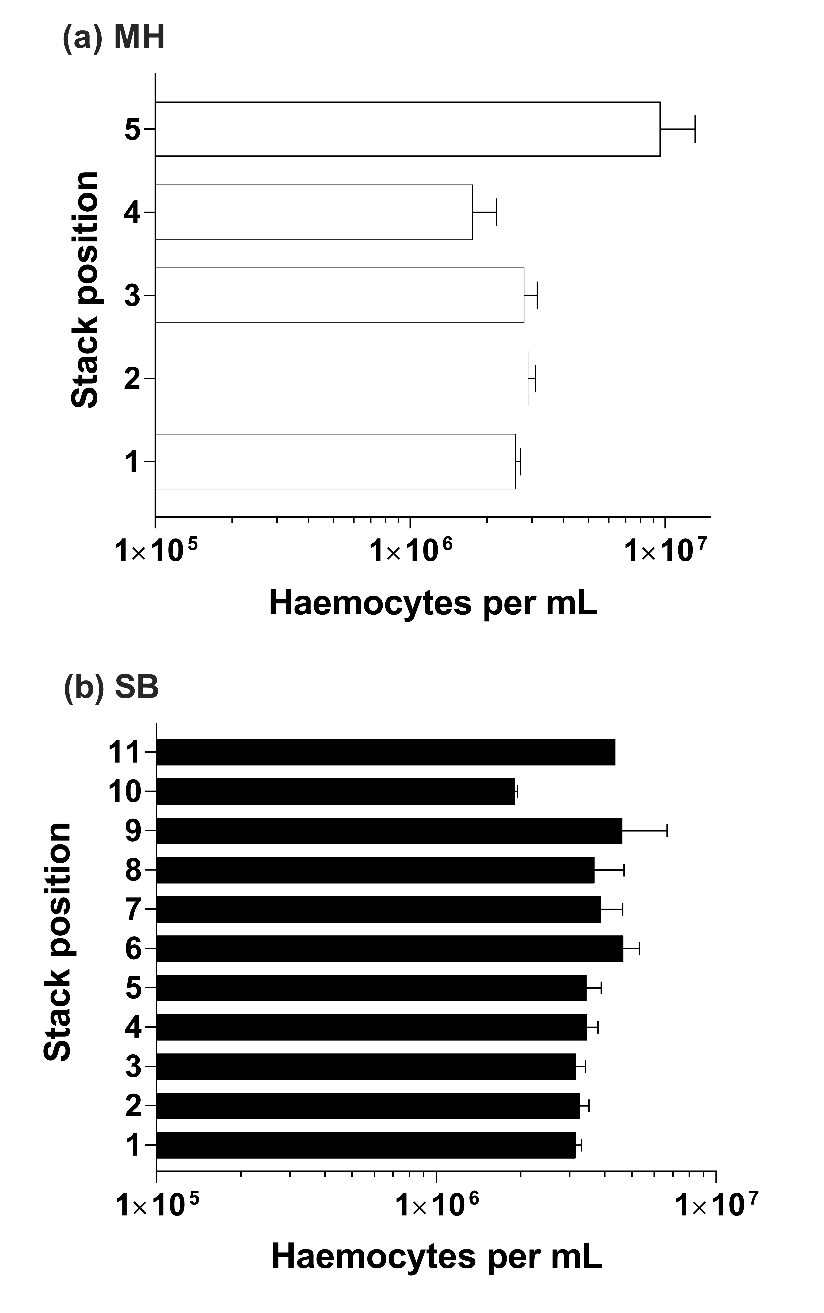

**Supplementary Figure 1. Total haemocyte counts in limpet haemolymph categorised by site and position within stacks.**

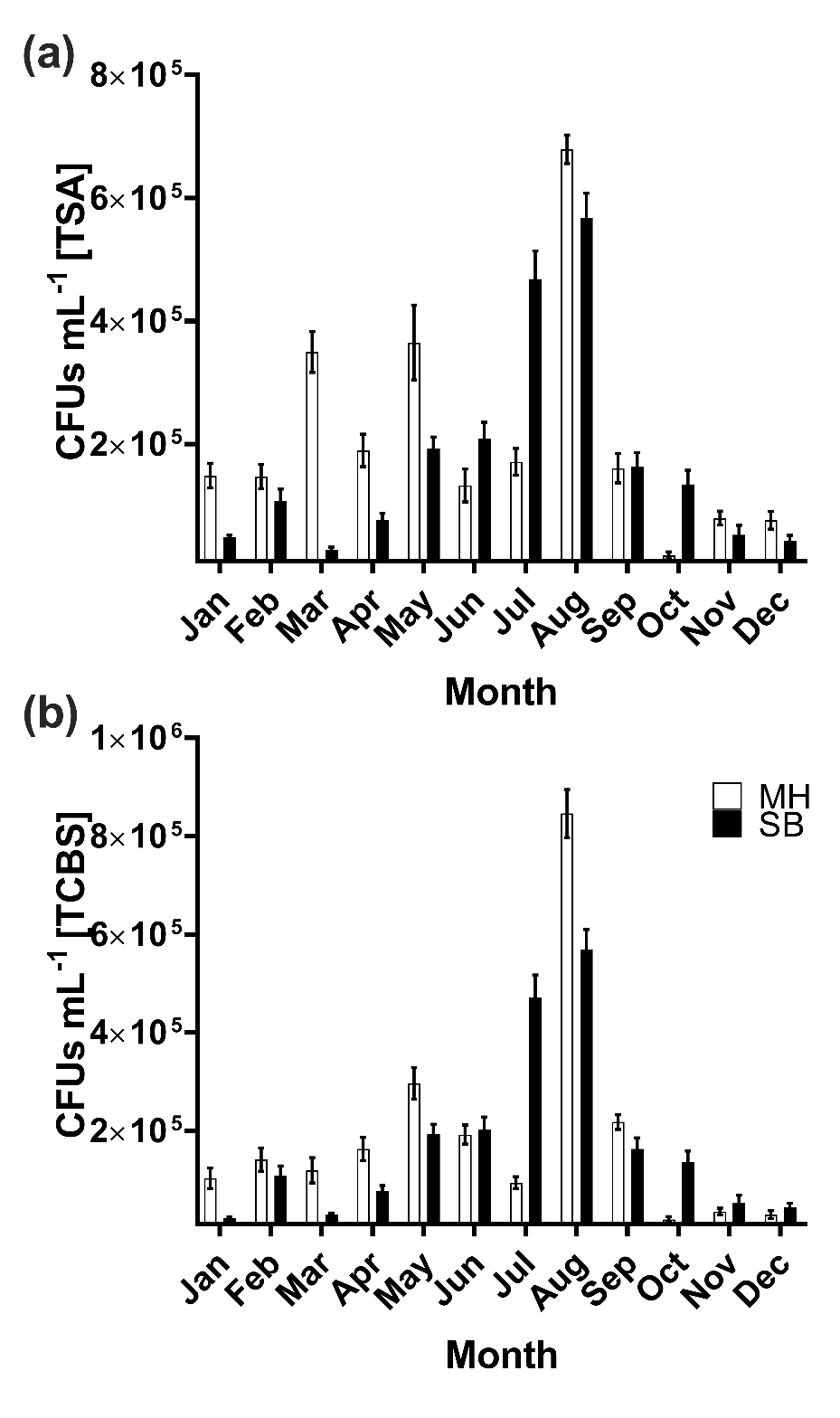

**Supplementary Figure 2. Abundance of bacterial colony forming units in limpet haemolymph enumerated using differential growth media.** Data are categorised according to site and month.

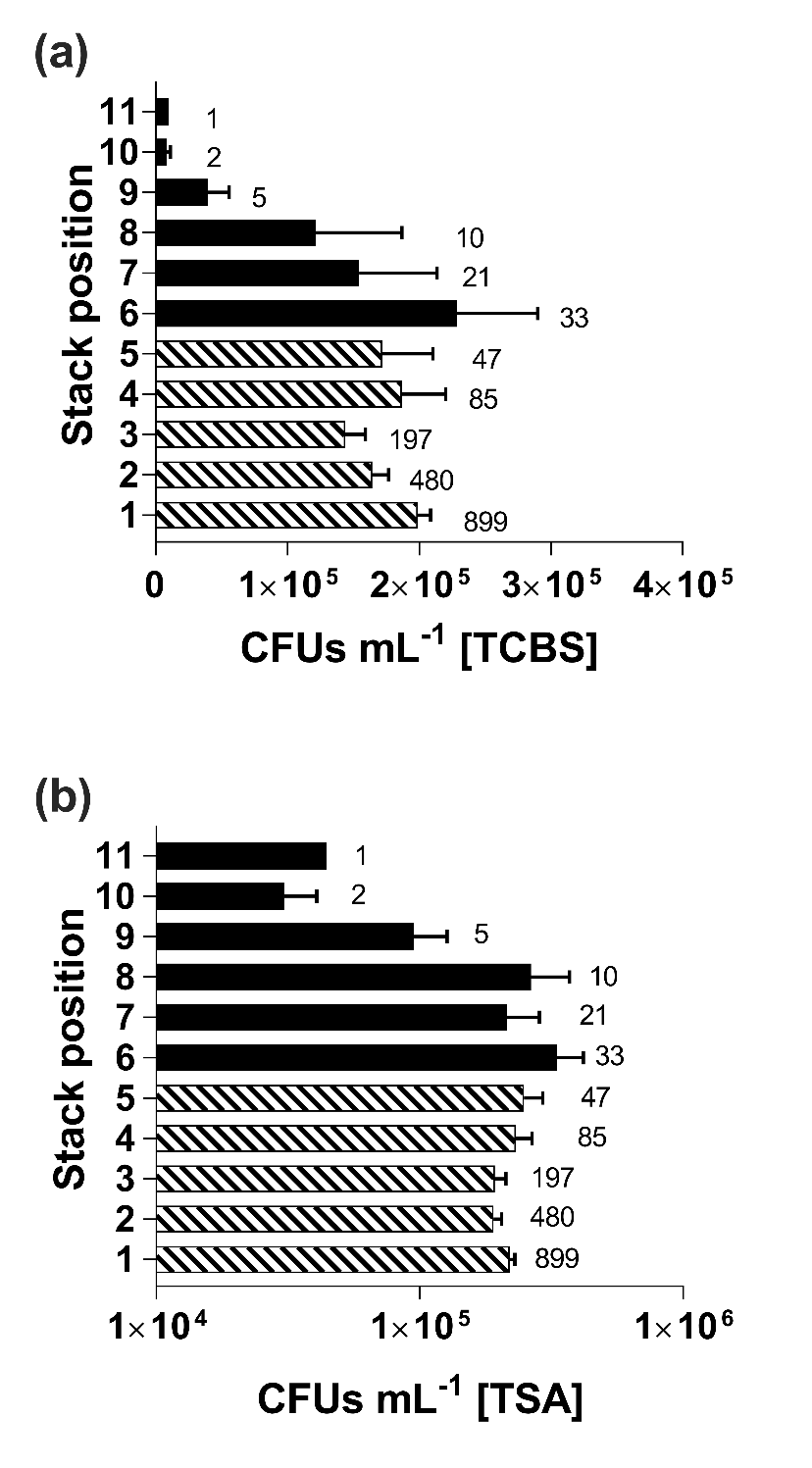

**Supplementary Figure 3. Abundance of bacterial colony forming units in limpet haemolymph enumerated using differential growth media.** Data are categorised according to position within stacks from both sites combined.

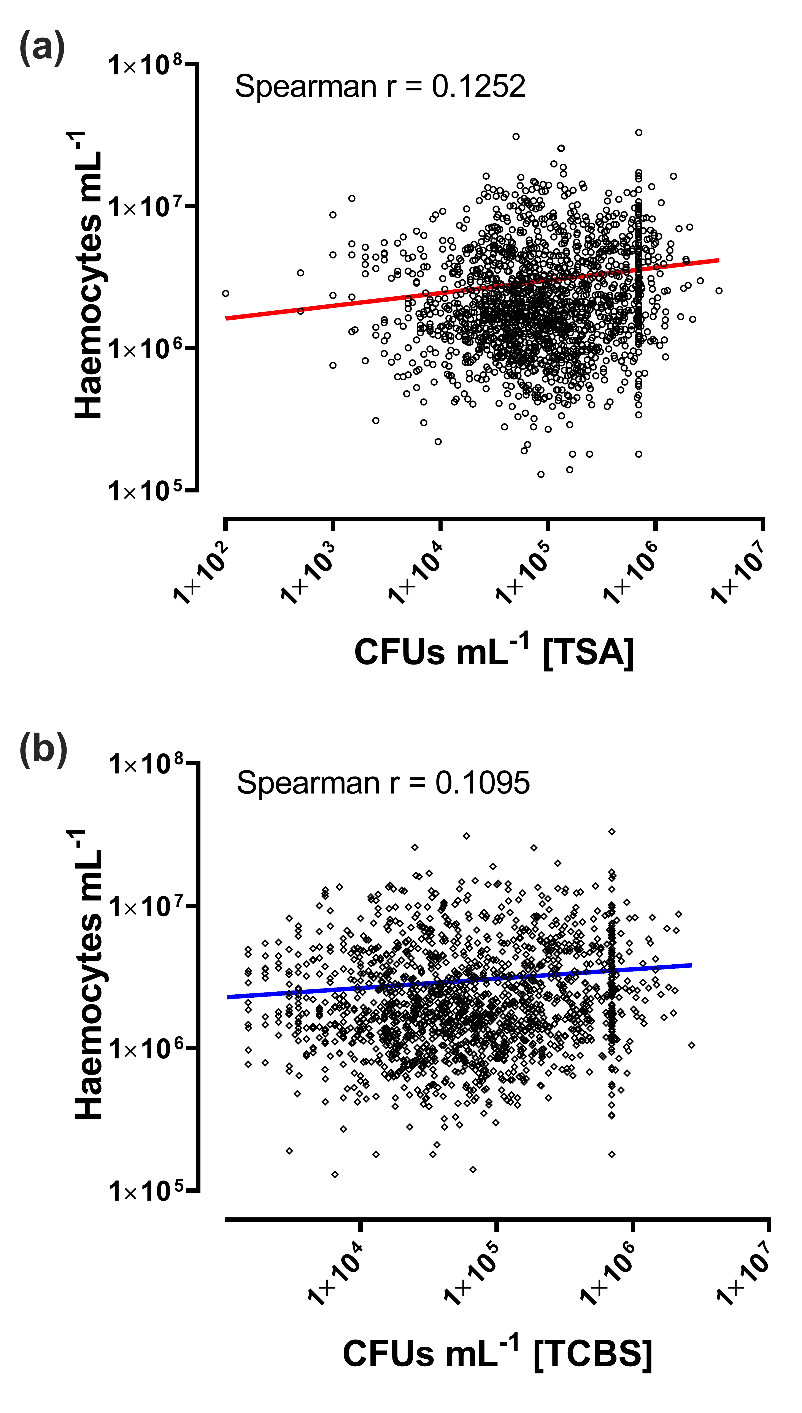

**Supplementary Figure 4 Relationship between bacterial colony forming units (CFU/mL) in the haemolymph of *C. fornicata* and observed haemocyte counts.** (a) *C. fornicata* haemolymph plated on generalist TSA. (a) Haemolymph plated on vibrio selective TCBS. Haemolymph bacterial load is expressed as CFUs per mL haemolymph (n = 1,770).

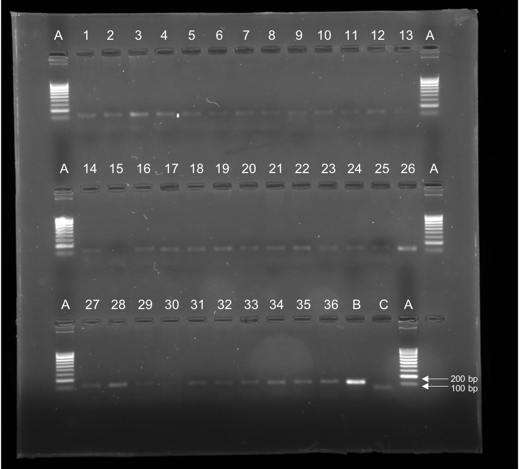

**Supplementary Figure 5. Agarose gel electrophoresis of PCR amplified products targeting vibrio sequences (16s V6 hypervariable region).**

100 bp

1000 bp

A - ladder (NZYTech, Lisboa, Portugal),

B - positive control consisted of 1 μL DNA purified from the haemolymph from an infected donor crab

C - Negative control (DEPC-treated ultra-pure water).

Wells 1-14, 16-29, and 31-36 are positive bands indicating amplification of vibrio products in *C. fornicata* sampled from Milford Haven in November 2019.

**Supplementary Table 1** BLASTn outputs from the 22 vibrio-like sequences amplified via PCR from *C. fornicata* haemolymph.

| Site | Month | Individual | | Closest Match | Accession Number | | QC(%) | | % ID | | Amplicon  Size | SRA Submission ID |
| --- | --- | --- | --- | --- | --- | --- | --- | --- | --- | --- | --- | --- |
| Jan | MH | | 13 | Uncultured bacterium clone SanDiego_a2787 | | KF799598.1 | 72 | 97.78 | | 61 | | SRR13165046 |
|  |  | |  | Uncultured bacterium clone SanDiego_a3071 | | KF799514.1 | 72 | 97.78 | |  | |  |
| Feb | MH | | 11 | *Vibrio mediterranei* | | HE584789.1 | 96 | 95.92 | | 50 | | SRR13165045 |
|  |  | |  | *Vibrio* sp. LiUU-B-16 | | DQ068946.1 | 92 | 95.74 | |  | |  |
| Feb | MH | | 11 | *Vibrio* sp. IO3 | | HQ848040.1 | 82 | 100 | | 64 | | SRR13165034 |
|  |  | |  | Vibrionaceae bacterium DS3 | | EF584037.1 | 81 | 100 | |  | |  |
| Feb | MH | | 25 | Uncultured bacterium clone SanDiego_a2787 | | KF799598.1 | 70 | 97.8 | | 62 | | SRR13165034 |
|  |  | |  | Uncultured bacterium clone SanDiego_a3071 | | KF799514.1 | 70 | 97.8 | |  | |  |
| Feb | MH | | 25 | *Vibrio tasmaniensis* isolate *25* | | EF178480.1 | 85 | 98.15 | | 63 | | SRR13165030 |
|  |  | |  | *Vibrio toranzoniae* strain 2-2 | | MT510186.1 | 85 | 96.3 | |  | |  |
| Sep | MH | | 39 | *Vibrio toranzoniae* strain 2-2 | | MT510186.1 | 86 | 100 | | 47 | | SRR13165029 |
|  |  | |  | *Vibrio crassostreae* strain 1-15 | | MT510175.1 | 100 | 100 | |  | |  |
| Oct | MH | | 43 | Uncultured bacterium clone SanDiego_a2787 | | KF799598.1 | 63 | 97.67 | | 66 | | SRR13165028 |
|  |  | |  | Uncultured bacterium clone SanDiego_a3071 | | KF799514.1 | 63 | 97.67 | |  | |  |
| Oct | MH | | 47 | Uncultured bacterium clone SanDiego_a2787 | | KF799598.1 | 95 | 93.65 | | 66 | | SRR13165027 |
|  |  | |  | Uncultured bacterium clone SanDiego_a3071 | | KF799514.1 | 95 | 93.65 | |  | |  |
| Nov | MH | | 55 | Uncultured bacterium clone DLPYS_MD02_081 | | KC852641.1 | 100 | 97.73 | | 44 | | SRR13165026 |
|  |  | |  | Uncultured bacterium clone SanDiego_a3077 | | KF799522.1 | 100 | 95.45 | |  | |  |
| Dec | MH | | 17 | Uncultured bacterium clone SanDiego_a2787 | | KF799598.1 | 72 | 100 | | 62 | | SRR13165025 |
|  |  | |  | Uncultured bacterium clone SanDiego_a3071 | | KF799514.1 | 72 | 100 | |  | |  |
| Dec | MH | | 17 | *Vibrio tasmaniensis* isolate 25 | | EF178480.1 | 92.4 | 98.11 | | 53 | | SRR13165044 |
|  |  | |  | *Vibrio toranzoniae* strain 2-2 | | MT510186.1 | 87.8 | 100 | |  | |  |
| Dec | MH | | 42 | Uncultured bacterium clone SanDiego_a2787 | | KF799598.1 | 87 | 91.38 | | 62 | | SRR13165043 |
|  |  | |  | Uncultured bacterium clone SanDiego_a3071 | | KF799514.1 | 87 | 91.38 | |  | |  |
| Dec | MH | | 42 | *Vibrio celticus* strain Rd 8.15 | | NR_116066.1 | 90 | 100 | | 60 | | SRR13165042 |
|  |  | |  | *Vibrio* sp. IO3 | | HQ848040.1 | 93 | 98.21 | |  | |  |
| Jan | SB | | 21 | Uncultured bacterium clone SanDiego_a2787 | | KF799598.1 | 100 | 97.78 | | 44 | | SRR13165041 |
|  |  | |  | Uncultured bacterium clone SanDiego_a3071 | | KF799514.1 | 100 | 97.78 | |  | |  |
| Jan | SB | | 21 | *Vibrio kanaloae* strain DS1904-Y1124 | | MT269633.1 | 100 | 100 | | 49 | | SRR13165040 |
|  |  | |  | *Vibrio gallaecicus* strain DS1904-Y1112 | | MT269632.1 | 100 | 100 | |  | |  |
| Jan | SB | | 52 | Uncultured bacterium clone SanDiego_a2787 | | KF799598.1 | 91 | 100 | | 49 | | SRR13165039 |
|  |  | |  | Uncultured bacterium clone SanDiego_a3071 | | KF799514.1 | 91 | 100 | |  | |  |
| Jan | SB | | 52 | Bacterium ocme08sprp595 | | JQ657678.1 | 90 | 96.55 | | 64 | | SRR13165038 |
|  |  | |  | Bacterium fme08x95np453l | | JQ657514.1 | 90 | 96.55 | |  | |  |
| Jan | SB | | 70 | Uncultured bacterium clone SanDiego_a2787 | | KF799598.1 | 90 | 93.55 | | 66 | | SRR13165037 |
|  |  | |  | Uncultured bacterium clone SanDiego_a3071 | | KF799514.1 | 90 | 93.55 | |  | |  |
| Jan | SB | | 70 | Uncultured bacterium clone YE-DC-A15 | | DQ438329.1 | 96 | 96.49 | | 58 | | SRR13165036 |
|  |  | |  | Uncultured bacterium clone A347_NCI | | FJ456660.1 | 96 | 94.64 | |  | |  |
| Feb | SB | | 45 | Bacterium ocme08sprp595 | | JQ657678.1 | 82 | 94.55 | | 64 | | SRR13165035 |
|  |  | |  | Bacterium fme08x95np453l | | JQ657514.1 | 82 | 94.55 | |  | |  |
| Aug | SB | | 45 | Uncultured bacterium clone SanDiego_a2787 | | KF799598.1 | 89 | 97.73 | | 48 | | SRR13165033 |
|  |  | |  | Uncultured bacterium clone SanDiego_a3071 | | KF799514.1 | 89 | 97.73 | |  | |  |
| Nov | SB | | 52 | Uncultured bacterium clone SanDiego_a2787 | | KF799598.1 | 93 | 95.56 | | 47 | | SRR13165032 |
|  |  | |  | Uncultured bacterium clone SanDiego_a3071 | | KF799514.1 | 93 | 95.56 | |  | |  |
